# Establishment and characteristic of an orthotopic implantation model of human hepatocellular carcinoma with Luc-GFP-labeled in nude mice

**DOI:** 10.1101/462887

**Authors:** Jing-jing Zhang, Min-hua Xu, Jie Wang, Xiao-bao Jin, Yan Ma

## Abstract

**Aim:** To construct Luc-GFP-labeled human hepatocellular carcinoma (HCC) cell line with high metastatic potential. And to establish a spontaneous metastasis and conveniently monitored orthotopic model of hepatocellular carcinoma in nude mice. Methods: HCCLM3-Luc-GFP cell line stably expressing luciferase (Luc) and green fluorescent protein (GFP) was constructed by lentivirus transfection. The orthotopic xenograft model was established though cell suspension injection method and tumor fragment implanted method. The growth and metastasis of the tumors were observed by in vivo imaging and pathology. Results: HCCLM3-Luc-GFP, a highly metastatic HCC cell line with GFP expression and Luc activity, was obtained. The tumorigenic rates both of two approaches were 100%, but the lung metastatic rate was higher the former than the latter. Conclusion: The orthotopic model of highly metastatic and Luc-GFP-labeled HCC in nude mice was successfully established by above approaches, called as cell suspension injection method and tumor fragment implanted method, respectively. This study provides a new and effective means to monitor the growth of tumors in vivo and to evaluate the efficacy of anti-metastatic drugs against HCC.

## INRTODUCTION

Hepatocellular carcinoma (HCC), the most common form of primary liver cancer, is the sixth commonest and the third most fatal malignancy worldwide^1^. In China, the incidence of liver cancer is high. There are 350,000 new cases and 320,000 deaths per year from liver cancer. In urban and rural areas of China, the mortality rate of liver cancer has risen to the second and the first in malignant tumors, respectively. The invasion, metastasis and recurrence of HCC are the leading causes of death ^3^.

Therefore, it’s important to explore the mechanisms of invasion, metastasis and recurrence of HCC and to find effective drugs to inhibit the metastasis and recurrence of HCC. Moreover, the establishment of a spontaneous metastatic and observable human hepatoma nude mouse model provide the basis of studying the recurrence and metastasis of human liver cancer and estimating the efficacy of anti-hepatoma drugs. Liver Cancer Institute of Fudan University had established high metastatic human hepatoma cell lines such as MHCC97H, HCCLM3 and high metastatic human hepatoma nude mice model for the treatment of liver cancer which can provide an effective means to study the metastasis and recurrence of Hepatoma. However, these tumors couldn’t be monitored continuously and intuitively in vivo. The quantitative of tumor metastasis is not accurate enough ^4,5^.

Optical in vivo imaging, using a very sensitive optical detection instrument, has become quantifiable, highly sensitive to monitor cell activity and gene behavior in living organisms, are being applied in pre-clinical research studies. Optical imaging is divided into fluorescence and bioluminescence. Fluorescence imaging works on the basis of fluorochromes inside the subject that are excited by an external light source, and which emit light of a different wavelength in response. Traditional fluorochromes include GFP, RFP, and their many mutants. Fluorescent luminescence is low-cost and simple^6,7^. However significant challenges emerge in vivo due to the autofluorescence of tissue. Bioluminescence imaging, on the other hand, is based on visible light generated by the oxidation of enzyme-specific substrates such as d-luciferin for terrestrial organisms and coelenterazine for marine organisms. Bioluminescence imaging does not require excitation of the reporter and has almost no background noise in vivo ^8,9^.

In this study, a lentiviral expression vector pCDH-CMV-Luc-EF1-GFP-Puro containing luciferase (Luc) gene and green fluorescent protein (GFP) gene, and human hepatocellular carcinoma cell line HCCLM3-Luc-GFP expressing the luciferase (Luc) and green fluorescence (GFP) were constructed. The orthotopic model of highly metastatic and Luc-GFP-labeled hepatocellular carcinoma in nude mice was established by cell suspension injection method and tumor fragment implanted method.. Then we observed the growth and metastasis of tumor in the model nude mice by noninvasive bioluminescence tomography and fluorescence excitation imaging (Fluorescence molecular tomography) at the living level. This study provides a novel and useful method for monitoring the growth and the metastasis of the tumor at living level, assessing the therapeutic effect of anti-hepatoma drugs.

## METHODS

### Cell Lines and Animals

The human hepatocellular carcinoma cell line HCCLM3 was purchased from the China Type Culture Collection(WuHan).. Cells were maintained in Minimum Essential Medium (MEM, Gibco BRL, Rockville, MD, United States) with 10% fetal bovine serum (Life Technologies, Carlsbad, CA, United States), and incubated at 37 °C in humidified atmosphere containing 5% CO2. HCCLM3 cells stably expressing the luciferase gene were constructed.

Female athymic BALB/c nude mice (6 weeks of age) were purchased from the Guangdong Laboratory Animal Center Co. Ltd. (Guangdong, China) and maintained in a pathogen-free animal facility at the Laboratory Animal Research Centre of Guangdong Pharmaceutical University. All experiments were performed with humane care, and were approved by the Animal Care and Use Committee at Guangdong Pharmaceutical University.

### Luciferase (Luc) expression plasmid construction and transfection

The DNA of the Luc gene was obtained by polymerase chain reaction (PCR) amplification of the psiCHECK-2 vector with primers specific to Luc. The forward primer P1: 5’-TGC <u>TCT AGA</u> ATG GAA GAC GCC AAA AAC ATA AAGA-3’ and the reverse primer P2: 5’-CGC <u>GGA TCC</u> TTA CAC GGC GAT CTT TCC GCC CTTC-3’ (P1 contains (underlined) the *Xba* I restriction site and P2 contains (underlined) the *BamH* I restriction site). The PCR conditions was 98°C for 10 s and 68 °C for 2 min (30 cycles). Then the Luc gene was cloned into the lentivirus expression pCDH-CMV-MCS-EF1-GFP-T2A-Puro vector and the positive clones were obtained by transducing to the *E. coli* DH5α cells and growing on LB plates with 50 μg/ml ampicillin. The plasmid containing the Luc gene was verified by restriction enzyme digestion by *Xba* I and *BamH* I and DNA sequencing. The correct recombinant plasmid pCDH-CMV-Luc-EF1-GFP-Puro was packed into lentivirus by Guangzhou Sagene Biotechnology Co., Ltd. (Guangzhou, China).

### Animal model

The HCCLM3 cells transfected with pCDH-CMV-Luc-EF1-GFP-Puro were digested by 0.25% pancreatin which in logarithmic growth period and resuspended by the MEM, and adjusted the concentration of the cells to 5×10^7^/ml for cell suspension injected method and 2×10^7^/ml for tumor fragment implanting method.

#### Cell-suspension injected method

The 6 to 8-week old female BLAB/c nude mice were anaesthetized by the pentobarbital sodium (50 mg/kg) and its abdomen were opened to expose the liver. Then the HCCLM3-Luc-GFP cells were injected slowly into the liver of nude mice by using the 28G aseptic micro injector, the liver was sent back to the abdomen and the abdominal wall was sutured. After the operation, the mice were sent back into the cages and raised in the SPF condition and the mice postoperative situation was observed by every day. The all experimental procedures used in this method should be operated under the sterile condition.

#### Tumor fragment implanting method

The HCCLM3-Luc-GFP cells were injected into right flank of the mice. Once the subcutaneous tumor reached a size of 1 to 1.5 cm in diameter, the mouse was sacrificed by the pentobar bital sodium (50 mg/kg). The tumor was resected under aseptic conditions and was cut into about 1 mm^3^ sections which were implanted into the left liver lobe of the nude mice then the mice were sent back into the cages and raised in the SPF condition and the mice postoperative situation were observed by every day. The all experimental procedures used in this method should also be operated under the sterile condition.

### Observation of the tumor growth and metastatic in nude mice model by in vivo imaging

For bioluminescence imaging and GFP fluorescence imaging. Mice were anesthetized using the luciferase substrate D-luciferin at a dose of 150 mg/kg in pentobarbital sodium (50mg/kg) administered by intraperitoneal injection on days 7, 14, and 35 after orthotopic implantation of HCCLM3-Luc-GFP cell or tumors. Growth and metastatic of the tumor tissues were monitored on the Tanon 5200Multi imaging system (Tanon Science & Technology Co. Ltd.). In accordance with the comparison of the strength of the luminescence, then the different between two methods of modeling and luminescence were compared.

### The evaluation of the tumor morphology and lung metastasis by pathological method

The nude mice were deeply anesthetized after the in vivo imaging, and the abdominal cavity and chest of the mice was opened to observe the tumor morphology, growth and metastatic. The lung tissues of the nude mice were fixed by the Bouin’s fluid(75 ml 0.9-1.2% picric acid, 5ml glacial acetic acid and 25 ml 40% formaldehyde) for 24h, and soaked with anhydrous ethanol for 12 h. The liver tissues and the lung tissues of the nude mice were processed using standard histological techniques, paraffin-embedded and stained with haematoxylin and eosin. The tumor growth and lung metastasis were evaluated.

## RESULTS

### The PCR amplification of luciferase (Luc) gene and the construction and identification of the recombinant expression vector

The Luciferase (Luc) gene was cloned from the psiCHECK-2 vector as template. The electrophoresis (Fig.1) showed that the length of the Luc gene was 1647bp, which was identical to the expected length. After the Luc was inserted into the pCDH-CMV-MCS-EF1-GFP-Puro vector (8189bp), the recombinant plasmid was identified by restriction enzyme digestion and sequence analysis (Fig 1). The plasmid was digested into two fragments which one was the Luc gene, and the other was the pCDH-CMV-MCS-EF1-GFP-Puro vector. These results indicated that the recombinant expression vector pCDH-CMV-Luc-EF1-GFP-Puro was constructed successfully.

**Fig. 1.**
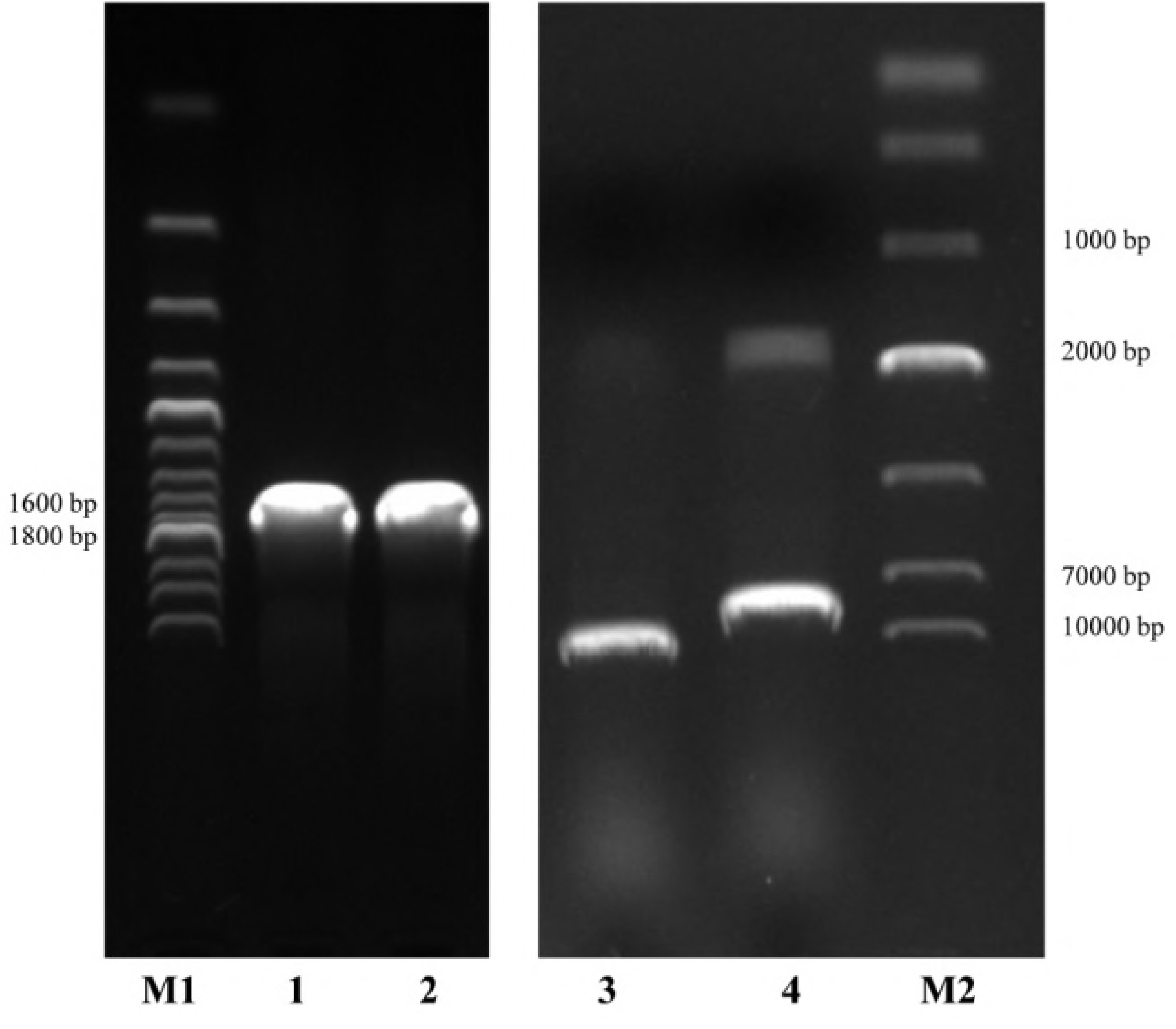
The electrophoretic results of the PCR amplification of luciferase gene and restriction map of recombinant expression vector. Lane M1 DNA Ladder (200bp), Lane M2 DNA Marker DL10000, Lanes 1–2 Luc gene fragments by PCR, Lane 3 pCDH-CMV-Luc-EF1-GFP-Puro plasmid digested with *BamH* I, Lane 4 pCDH-CMV-Luc-EF1-GFP-Puro plasmid digested with *Xba* I and *BamH* I.

### The construction and identification of the HCCLM3-Luc-GFP cell lines with GFP expression and luciferase activity

The HCCLM-3 cells were observed by inverted fluorescence microscope after the transfection with lentivirus for 72h. Compared with non-transfected cells, transfected cells were no distinct differences in cell morphology. On the contrary, the transfected cells had a striking increase in GFP fluorescence compared to the non-transfected cells (Fig 2). Moreover, the Luciferase activity in the transfected group was higher than in the non-transfected group, and the value in the transfected group increased with increasing concentration (Tab 1). These results indicated that the HCCLM3-Luc-GFP cell lines with GFP expression and luciferase activity was constructed successfully.

**Table. 1.** Luciferase activity assay in HCCLM3-Luc-GFP cells.

| Concentration | Non-transfected | Transfected |
| --- | --- | --- |
| 8*10<sup>5</sup> | 20.22 | 95827.25 |
| 4*10<sup>5</sup> | 20.56 | 45955.97 |
| 2*10<sup>5</sup> | 21.56 | 12801.60 |
| 1*10<sup>5</sup> | 22.38 | 5543.15 |

**Fig. 2.**
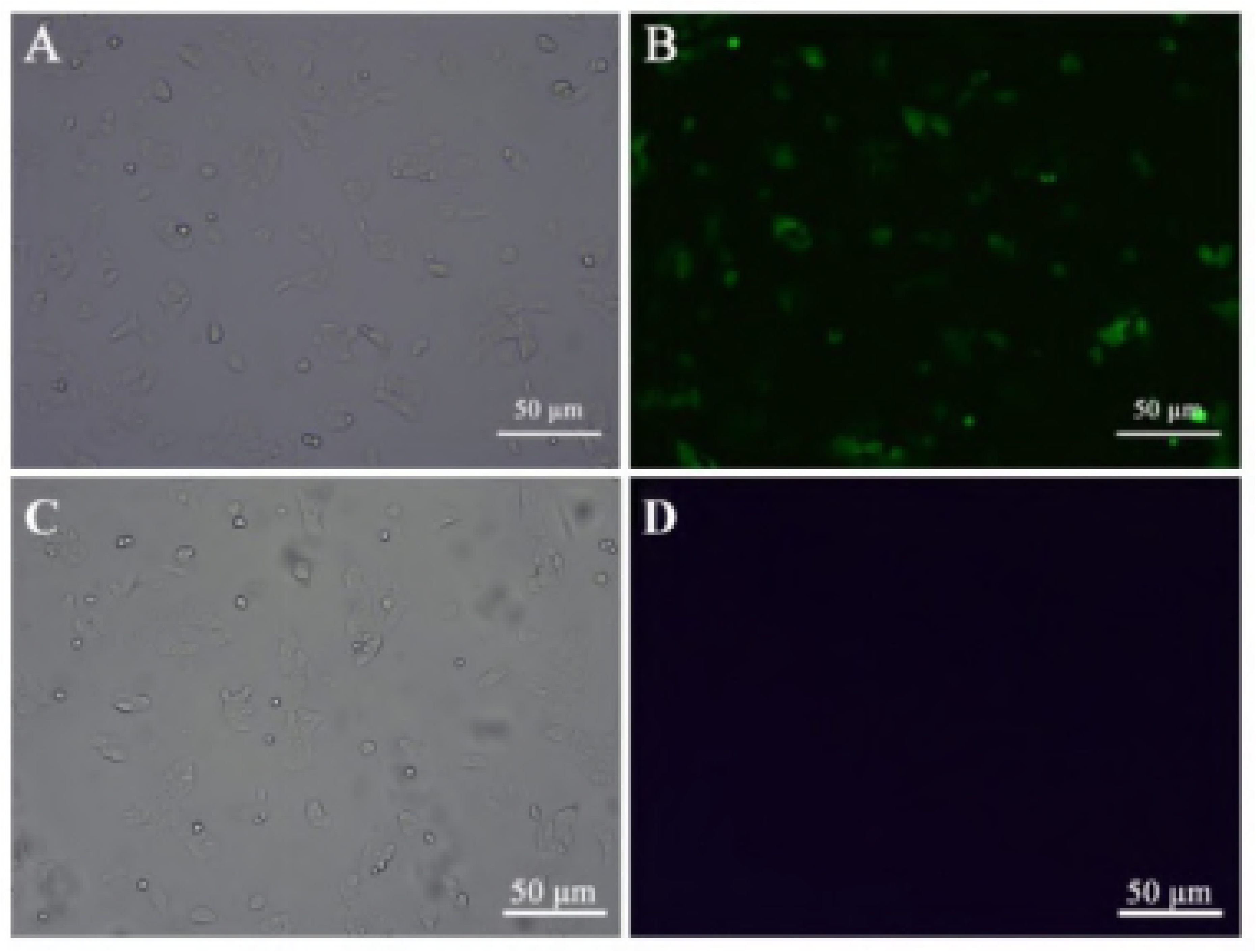
The GFP observation of transfected (A, B) and non-transfected (C, D) cells under the white light (A, C) and fluorescence microscopy (B, D).

**Fig. 3.**
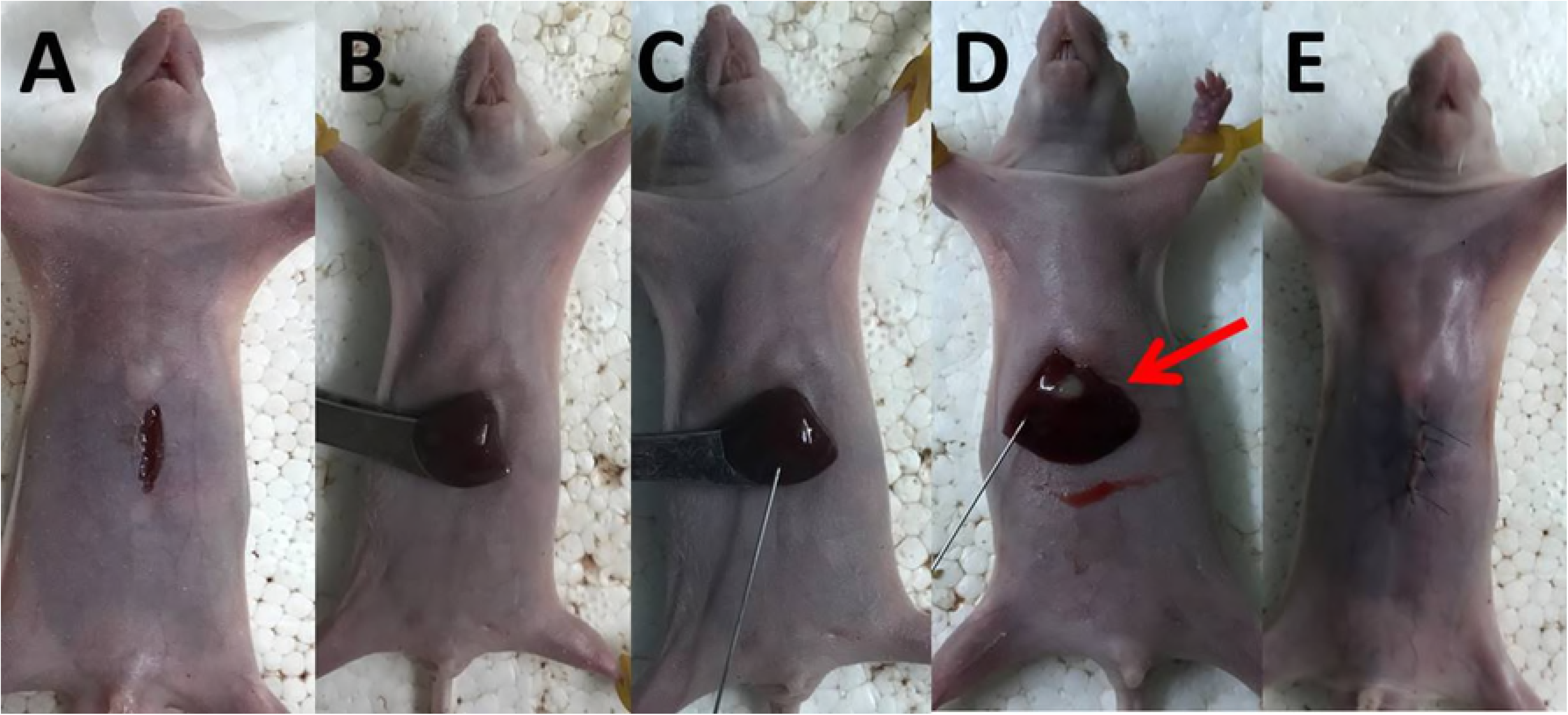
The operation process of the model constructed by cell suspension injected method.

**Fig. 4.**
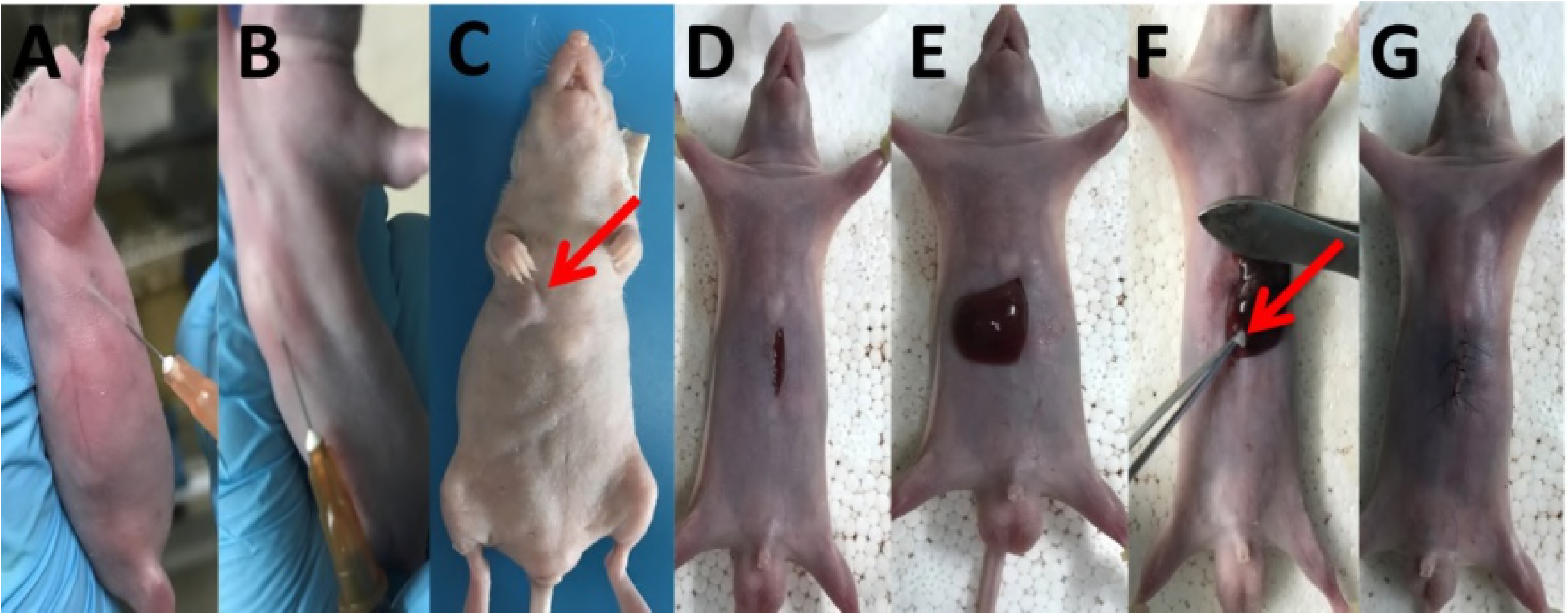
The operation process of the model constructed by tumor fragment implanting method.

### The in vivo imaging observation of the nude mouse model constructed by cell suspension injected method and tumor fragment implanting method

After liver orthotopic xenograft, the bioluminescence signal and GFP fluorescence signal of the model was observed by the Tanon 5200Multi animals in vivo imaging system. The bioluminescent signal of cell suspension injected method can be observed for 2 weeks (Inoculation amount was10μl, Fig 5A_1_) or 1 week (Inoculation amount was 20μl, Fig 5A_2_). Moreover, the tumor fragment implanting method was observed at 2 weeks. The signal of the GFP luminescence can be seen at 3 weeks and is easily interfered by outside situation (Fig 5B). However, the GFP luminescence signal of tumor fragment implanting method could not be observed (Fig 6). The results also demonstrated that the bioluminescent signal and GFP fluorescence signal increased with the increase of the irradiation time and the fluorescence would reach the highest after 5 weeks.

**Fig. 5.**
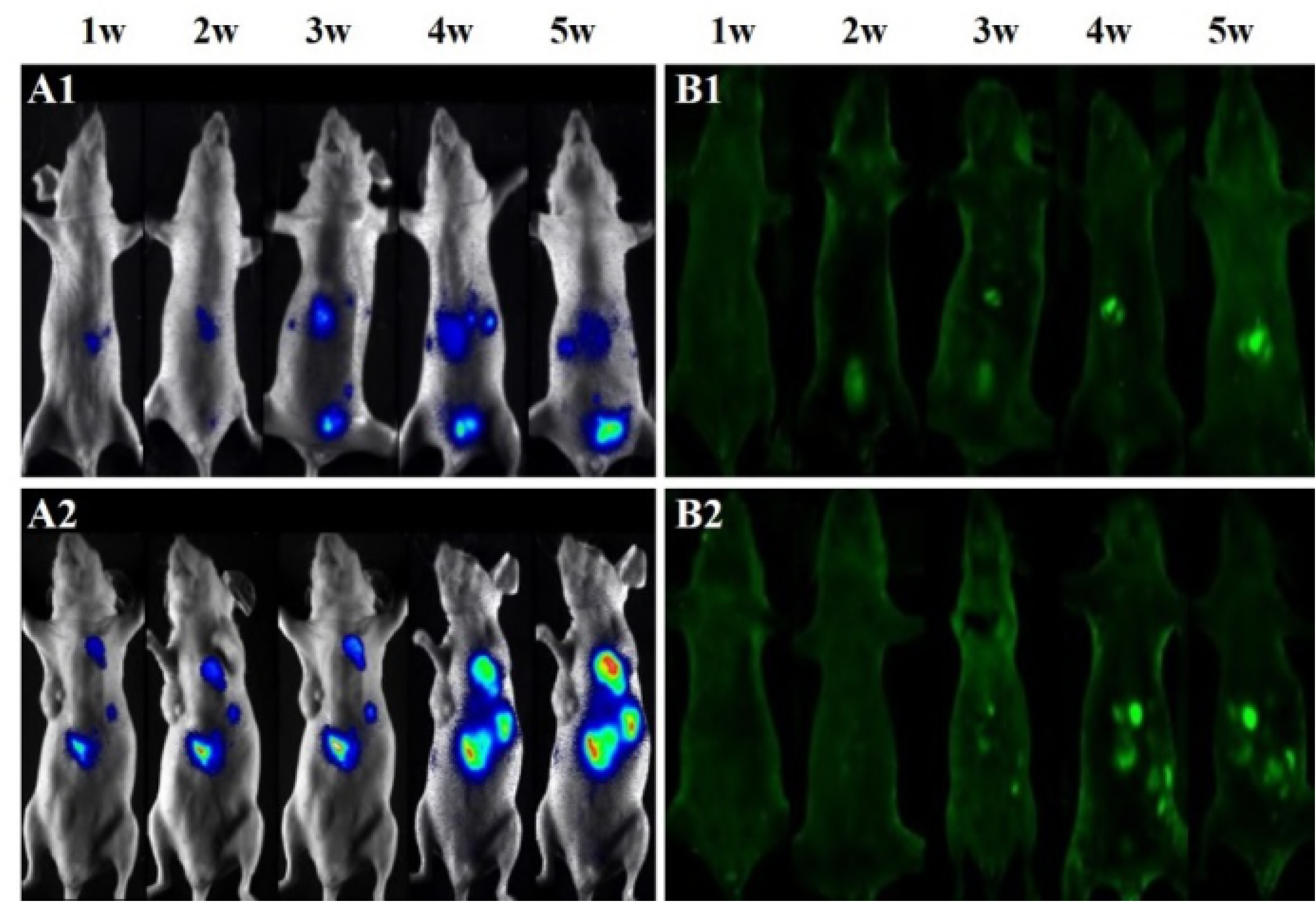
In Vivo Imaging of the orthotopic xenograft model constructed by cell suspension injected method. (A) Under the Luc bioluminescence detection. (B) Under the GFP green fluorescence detection. (1) Inoculation amount was10μl. (2) Inoculation amount was 20μl

**Fig. 6.**
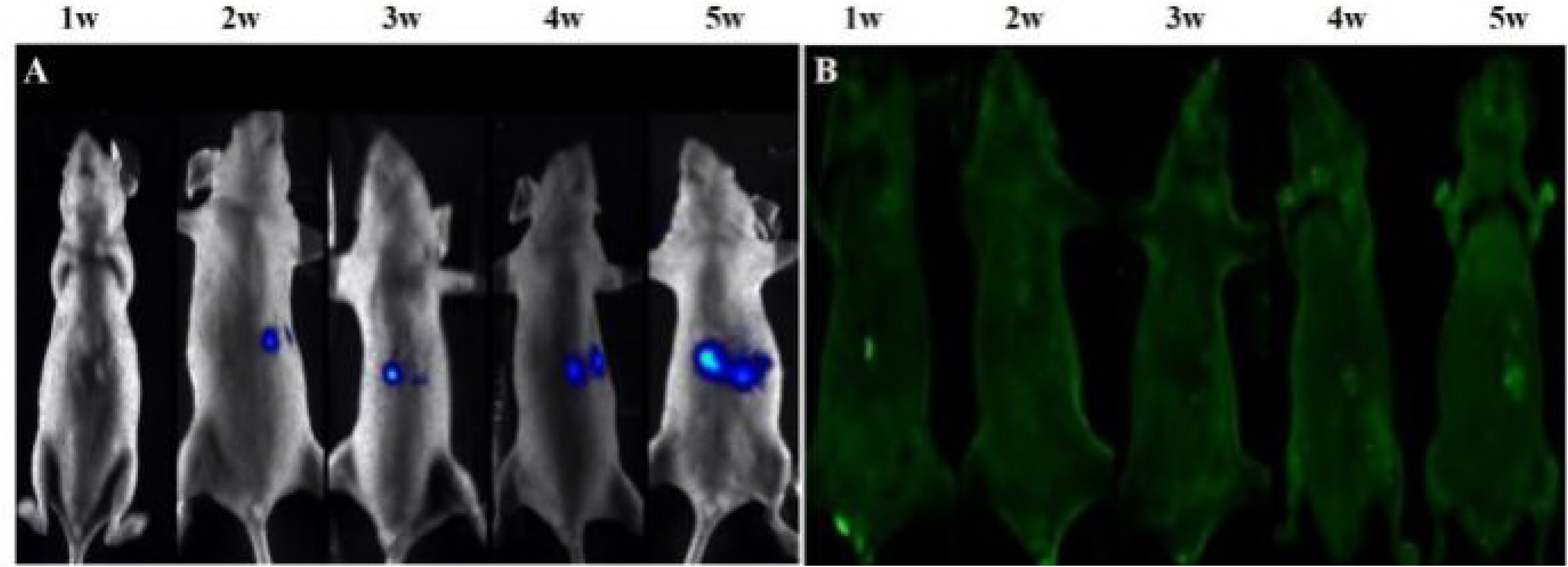
In Vivo Imaging of the orthotopic xenograft model constructed by tumor fragment implanting method. (A) Under the Luc bioluminescence detection. (B) Under the GFP green fluorescence detection.

### The morphological observation of the tumor tissues

The nude mice were dissected to obtain the tumor tissue after the growth of the bearing-mice for 5 weeks. There are multiple tumor nodules on the liver surface in the mice by the cell suspension injected method, and they were well-distributed on the liver (Fig 7A). The tumor tissues in the mice by tumor fragment implanting method were distributed on the surface and the edge of the liver tissues and could not permeate the whole liver (Fig 7B).

After the lungs are fixed, the metastases are light yellow or white, and the lungs are yellow-brown. The model of cell suspension injected method can be seen as a gross lung metastasis with single or multiple isolated nodules. (Fig 7C.) The lung tissue collapsed and no gross metastases by tumor fragment implanting method. (Fig 7D.) Moreover, HE staining further confirmed tumorigenesis of the liver tissue (Fig 8)and the lung metastasis. (Fig 9)

**Fig. 7.**
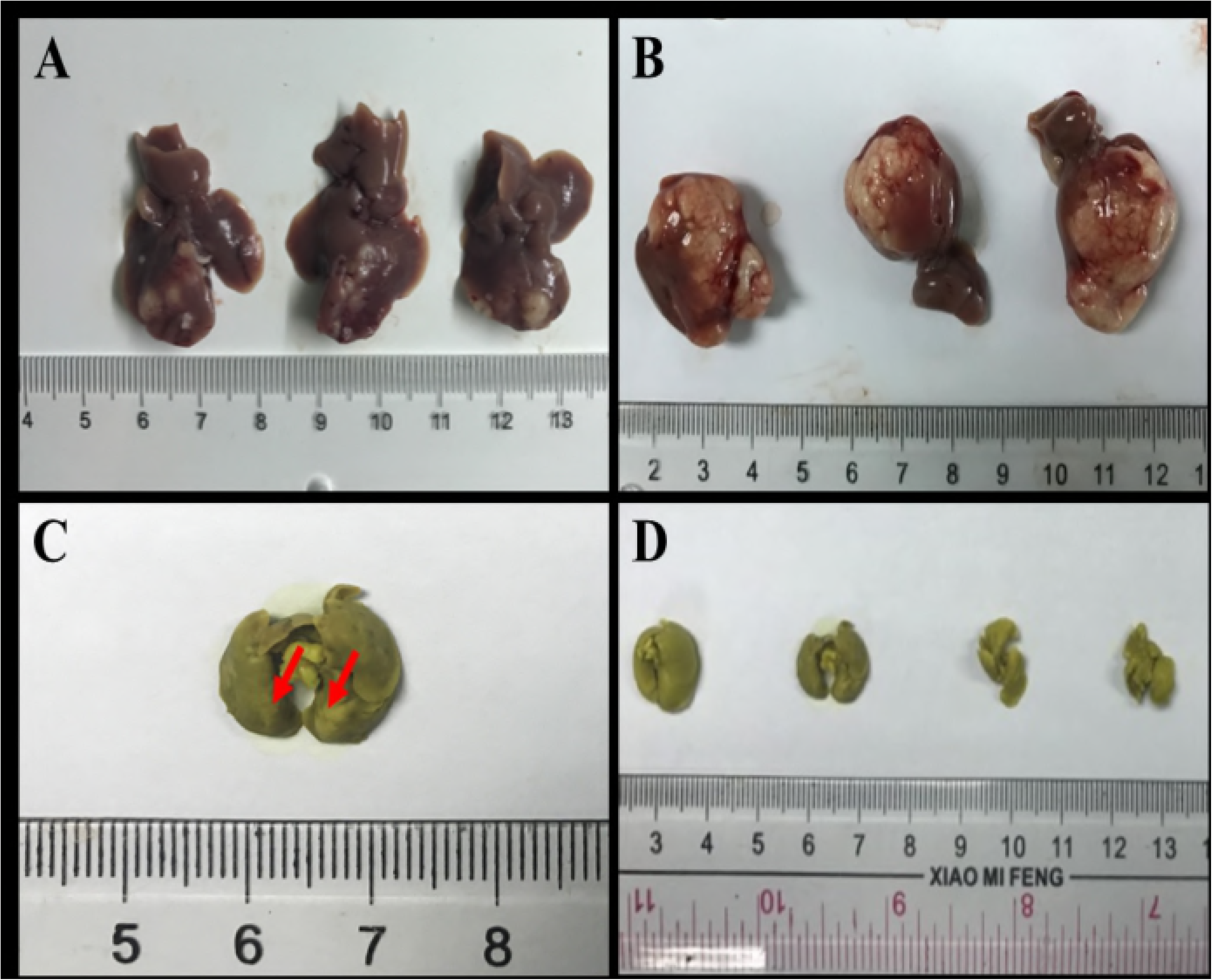
The physical observation of the liver tumor tissues (A, B) and the lung tumor tissues (C, D) in the orthotopic xenograft model constructed by cell suspension injected method (A, C) and tumor fragment implanting method (B, D)

**Fig. 8.**
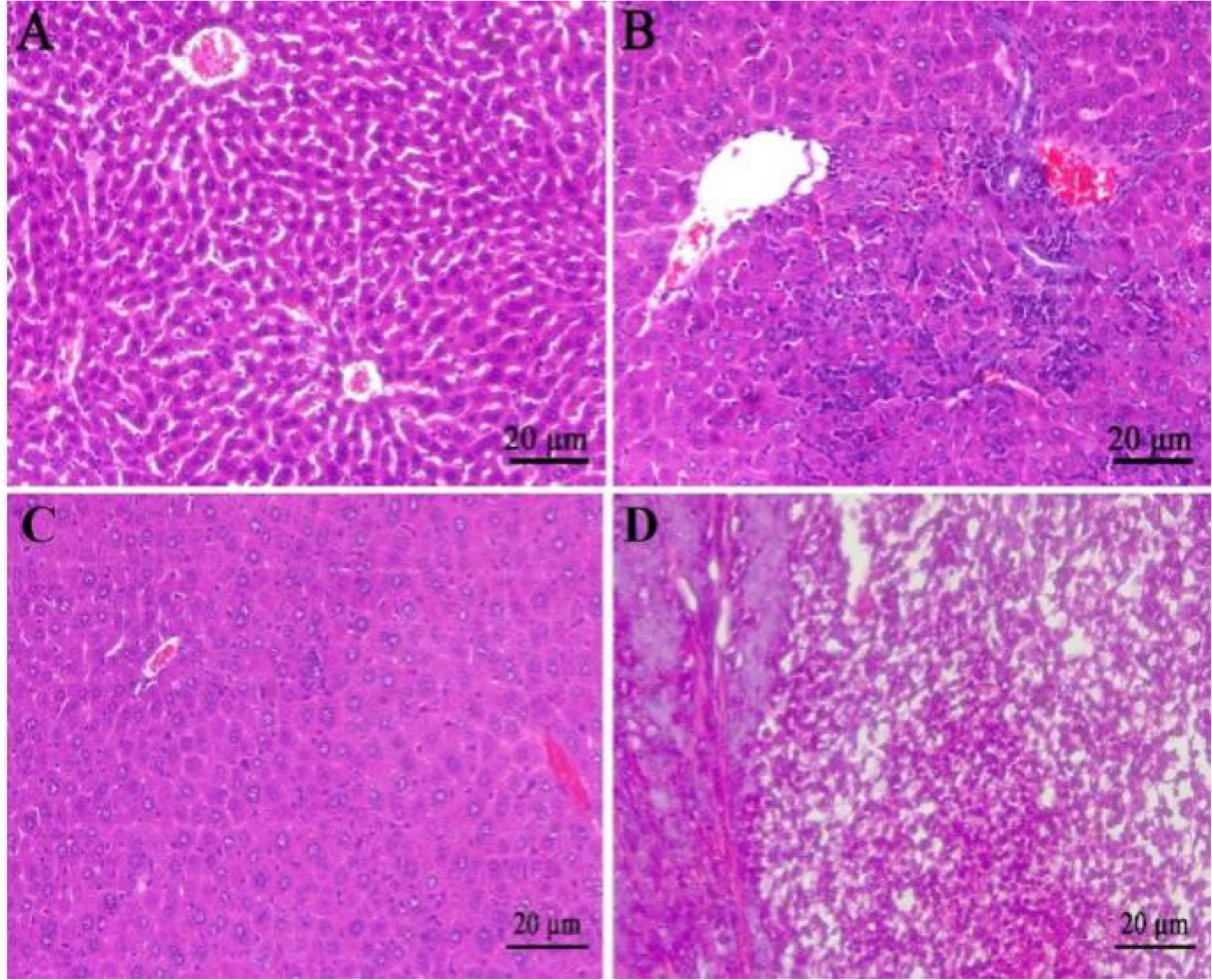
The HE staining results of the normal liver tissues (A, C) and the liver tumor tissues(B, D) in the orthotopic xenograft model constructed by cell suspension injected method (A-B) and tumor fragment implanting method (C-D)

**Fig. 9.**
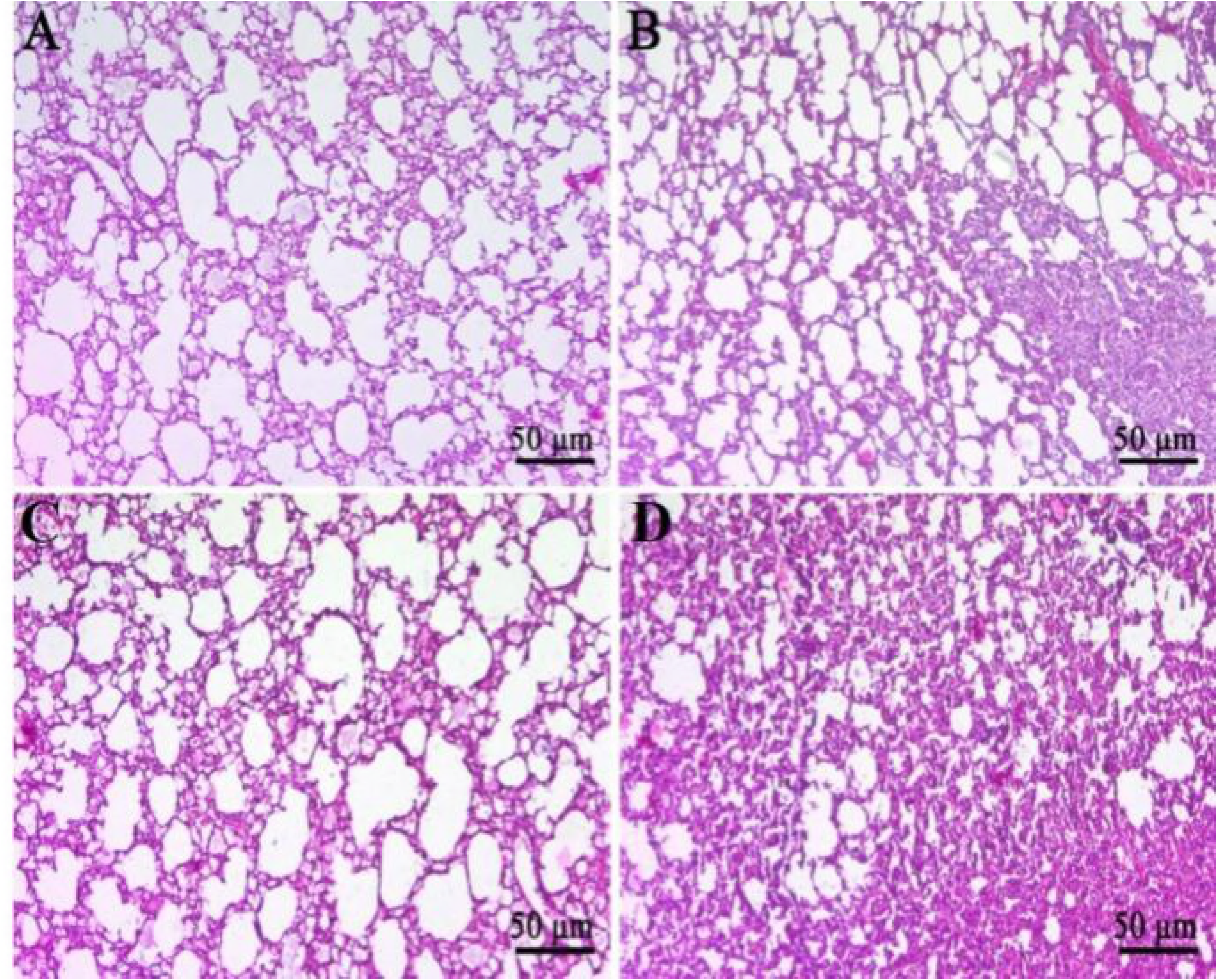
The HE staining results of the normal lung tissues (A, C) and the lung tumor tissues (B, D) in the orthotopic xenograft model constructed by cell suspension injected method (A-B) and tumor fragment implanting method (C-D)

## DISCUSSION

The nude mouse orthotopic implantation model of human hepatocellular carcinoma is an important tool to study the underlying mechanism of the occurrence and metastasis of liver cancer, and it is a new approach to explore anti-hepatoma drugs ^10,11^. At present, this model is mostly constructed by tumor fragment implanted and cell suspension injection^12^, the two methods have their own characteristics, but there are shortcomings. Tumor fragment implanted method, the goal is to establish a subcutaneous model of human HCC in nude mice, and then to inoculate the liver in order to establish the orthotopic implantation model. The survival rate of the implanted tumor is supposed to increase. Use of this model is limited due to the need for a highly complex surgical technique, especially in bleeding control. Compared with tumor fragment implanted method, the invasion and metastasis of the orthotopic tumor model by cell suspension injection method are better. However, the technical difficulty associated with this model is that it is hard to control tumor cell leakage during injection^13^. Analyses are complicated by the fact that apparent metastasis may just be due to leakiness. In this paper, the nude mouse orthotopic implantation model of human hepatocellular carcinoma were successfully established by the two different methods, and their tumorigenic rate were 100%. The former takes longer time, and the tumor grow in the implant site of liver. The latter can be shortened by half the time and the tumor can form in whole liver. The micro-environment of the latter is more relevant to the process of tumor growth in clinical practice; thus, it can be a useful research model for study the metastasis and recurrence of liver cancer. The results of this study are consistent with the descriptions reported in the above literature.

To explore the mechanism of invasion and metastasis of hepatoma and to seek drugs which can inhibit the invasion and metastasis of liver cancer, a conveniently monitored orthotopic model of hepatocellular carcinoma in nude mice is essential. As a photoprotein, green fluorescent protein (GFP) has been widely used in the intracellular molecular imaging because of many advantages such as no need for substrate, low cost and simple operation. However, the excitation and emission wavelengths of GFP are short and they are easily limited by the interference of external substances and the permeability of the spectrum. Therefore, GFP has a high background or is not easily detected in vivo imaging^14^. Luciferases (Luc) is a generic term of oxidative enzymes that produce bioluminescence, and is usually distinguished from a photoprotein. Luc has no endogenous expression in mammals and it is not affected by other substances, so it can accurately monitor the primary tumor and the tiny metastases tumor. As a reporter gene, luciferase provides an accurate and dynamic monitoring means for early tumor in vivo ^15-17^. However, luciferase requires addition of luciferin, the consumable substrate, which is complicated and expensive to perform. In this study, green fluorescent protein and luciferase, as reporter genes, were used for molecular imaging, which combined the advantages of both while can compensate for the deficiencies of the other^18-20^. The results showed that bioluminescence signal was detected by in vivo imaging on the first day after inoculation of HCCLM3-Luc-GFP cells in the liver of nude mice. As time goes by, the fluorescence signal enhanced gradually, with high sensitivity and specificity. However, GFP showed weak luminescence in subcutaneous and liver in situ, and almost no significant change of the fluorescence signal was detected. In the nude mouse model of hepatocellular carcinoma, the bioluminescence effect of Luc was significantly better than that of GFP-labeled fluorescence imaging.

In order to establish a spontaneous metastatic hepatocellular carcinoma xenograft model, HCCLM3, high metastatic human hepatocellular carcinoma cells, was selected as the research object in this study. The orthotopic model of highly metastatic hepatocellular carcinoma in nude mice was established by tumor fragment implanted method and cell suspension injection method,. The tumor and HE staining showed that the liver tumor in situ which was established by the former method only grew in the implant site, and could not transfer to other parts of the liver. The liver tumor in situ which was established by the latter method was detected for intrahepatic and abdominal metastases and the rate of lung metastasis was higher than the former. In addition, we also observed a high abdominal metastases in cell suspension injection group. One of the reasons for this high metastases may have been due to the tumor cell leakage. Because the fluorescence intensity of the abdomen in models, which were injected 20ul human HCC cells, was stronger than that in models with 10ul. Under the premise of being able to build the models, we would like to point out that the smaller volume of the HCC cells was better. In this study,a cotton swab was used to prevent leakage during injection. Cells may leak into the abdominal cavity becauese of post-operative activities. Therefore, we would optimize the method to seal the wound and reduce the cells into the abdominal cavity in the later study^21^.

In summary, we investigated the tumor growth and metastasis of HCC in nude mice established by cell suspension injection method and tumor fragment implanted methodc. We also compared the imaging effect of luciferase and GFP in the nude mouse with orthotopic liver cancer model. This study will be useful for establishing a conveniently monitored and spontaneous metastasis orthotopic liver cancer model to evaluate the efficacy of anti-metastatic drugs against hepatocellular carcinoma.

## ACKNOWLEGEMENTS

This work was supported by National Natural Science Foundation of China (NO. 81502520), Natural Science Foundation of Guangdong Province of China (NO. 2016A030310299) and Medical Scientific Research Foundation of Guangdong Province of China (NO. A2016091).

## REFERENCES

1. Ahmedin J, Freddie B, Melissa M.C,et al. Global Cancer Statistics. CA Cancer J Clin, 2011, 261(2):69–90

2. Gao J, Xie L, Chen WQ, et al. Rural-urban, sex variations, and time trend ofprimary liver cancer incidence in China. Eur J Cancer Prev, 2013, 22(5):448–54

3. Liover JM, Burroughs A, Bruix J, et al. Hepatocellular carcinoma. Lancet, 2003, 362(9399):1907–1917

4. Li Y, Tang ZY, Ye SL, et al. Establishment of cell clones with different metastatic potential from the metastatic hepatocellular carcinoma cell line MHCC97. World J Gastroenterol, 2001, 7(5):630–636

5. Li Y, Tian B, Yang J, et al. Step wise metastatic human hepatocellular carcinoma cell model system with multiple metastatic potentials established through consecutive in vivo selection and studies on metastatic characteristics. Joumal of Cancer Research and Clinical oncology, 2004, 130(8):460–480

6. Yang BW, Liang Y, Xia JL, et al. Biological characteristics of fluorescent protein-expressing human hepatocellullar carcinoma xenograft model in nude mice.European Journal of Gastroenterology & Hepatology, 2008, 20(11):1077–84

7. Chen Q, Wang XP, Wu H, et al. Establishment of a dual-color fluorescence tracing orthotopic transplantation model of hepatocellular carcinoma. MOLECΜLAR MEDICINE REPORTS, 2016, 13(1):762–768

8. Wang Q, Luan W, Vadim Goz, et al. Non-invasive in vivo imaging for liver tumor progression using an orthotopic hepatocellular carcinoma model in immunocompetent mice. Liver Internationa, 2011, 31(8):1200–8

9. WuT, Heuillard E, Lindner V, et al. Multimodal imaging of a humanized orthotopic model of hepatocellular carcinoma in immunodeficient mice. Scientific Reports, 2016, 14(6):35230–35230

10. Huynh H, Ngo VC, Koong HN, et al. AZD6244 enhances the anti-tumor activity of sorafenib in ectopic and orthotopic models of human hepatocellular carcinoma (HCC). Journal of Hepatology, 2010, 52(1):79–87

11. Huynh H, Chow PK, Tai WM, et al. Dovitinib demonstrates antitumor and antimetastatic activities in xenograft models of hepatocellμLar carcinoma. J Hepatol, 2012, 56(3):595–601

12. Wang SJ, Wei AL, Zhang YQ, et al. Comparison on the Three Kinds of Orthotopically Transplanted Model of Liver Transplantation Carcinoma in Rats many human diseases. Laboratory Animal Science, 2012; 29: 11–14.

13. Song XF, Lv Z, Liu X, et al. Comparaison of Intrahepatic Injection and Tumor Tissue Transplantation to Create Hepatic Tumor Model in Rats. Chinese Journal of Comparative Medicine, 2007; 17: 96–98.

14. Dufort S, Sancey L., Wenk C, et al. Optical small animal imaging in the drug discovery process. Biochimicaet Biophysica Acta, 2010, 1798(12):2266–2273

15. Czupryna J, Tsourkas A, et al. Firefly luciferase and RLuc8 exhibit differential sensitivity to oxidative stress in apoptotic cells. PLoS 0ne, 2011, 6(5):e20073–e20073

16. Liang Y, Walczak P, BμLte J W, et al. Comparison of red-shifted firefly luciferase Ppy RE9 and conventional Luc2 as bioluminescence imaging reporter genes for in vivo imaging of stem cells. J Biomed Opt, 2012, 17(1):16004–16004

17. Che P, Cui L, Kutsch O, et al. Validating a firefly luciferase-based high-throughput screening assay for antimalarial drug discoveiy. Assay Drug Dev Technol, 2012, 10(1):61–68

18. Deluca M.Transient and stable expression of the firefly luciferase gene in plant cells and transgenic plants. Science, 1986, 234(4778):856–859

19. Michelini E, Cevenini L, Mezzanotte L, et al. Luminescent probes and visualization of bioluminescence. MethodsMol Biol, 2009, 574(574):1–13

20. Navizet I, Liu YJ, Ferre N, et al. The chemistty of biolu-minescence:an analysis of chemical functionalities. Chemphyschem, 2011, 12(17):3064–3076

21. Kollmar O,Schilling MK, Menger MD, et al. Experimental liver metastasis: Standards for local cell implantation to study isolated tumor growth in mice. Clinical & Experimental Metastasis, 2004, 21(5):453–460

